# Clinical *Trypanosoma cruzi* isolates share a common antigen repertoire that is absent from culture adapted strains

**DOI:** 10.1101/2025.06.04.657671

**Authors:** Jill M.C. Hakim, Sneider A.G. Guiterrez, Angel Duran, Edith Malaga-Machaca, Carolina Duque, Lulu Singer, Rony Colanzi, Jacqueline E. Sherbuk, Caryn Bern, Robert H. Gilman, Lousia A. Messenger, Monica R. Mugnier, Working Group on Chagas Disease in Bolivia and Peru

## Abstract

**Background:** *Trypanosoma cruzi* causes Chagas disease, a poorly understood and clinically heterogeneous disease. Recent work has demonstrated that parasites adapted to laboratory conditions are genomically variable, but little is known of the extent of genomic diversity from clinically isolated specimens.

**Methods:** In this retrospective observational genomic study, we isolated 15 *T. cruzi* specimens from three clinical studies of Chagas disease, representing different clinical contexts. We sequenced the genome of each strain and used single nucleotide variant (SNV) based analyses to estimate parasite genetic lineage, genomic population structure, regions of copy number plasticity, and to identify gene conversion events. In addition, we generated and annotated whole genome assemblies of each isolate. From these assemblies, we compared the repertoires of genes encoding for highly virulent and variable proteins that have been implicated in disease pathogenesis.

**Findings:** We identified parasites from two genetic lineages in this collection of clinical isolates. Our analysis revealed evidence of genomic instability. Diversity-generating copy number variation was statistically enriched in regions encoding the virulence-associated multigene families, while diversity-eliminating gene conversion events were enriched in regions depleted of multigene family members. We also discovered a set of multigene family members that is present in all of the clinically isolated parasite genomes and absent from all of the lab adapted strains, regardless of parasite lineage. Multigene family repertoires were more conserved among field isolated specimens of the same genetic lineage than among culture adapted strains of the same genetic type.

**Interpretation:** This study provides whole genome sequencing data for TcV parasites isolated from naturally infected human patients with Chagas disease for the first time. Our analysis of these genomes revealed substantial genomic instability, suggesting the parasite undergoes genomic change in response to the pressures imposed by the host environment. Moreover, we observed a set of virulence-associated genes that are present exclusively within clinical isolates and absent from lab-adapted strains, indicating a potential role for these genes in parasite survival in natural hosts. These findings highlight the limitations of genetic studies focused exclusively on lab-adapted parasite strains and provide insight into the genomic features of *T. cruzi* that are likely to be important for clinical infection.

## INTRODUCTION

*Trypanosoma cruzi*, the causative agent of Chagas disease, a neglected tropical disease, is currently estimated to infect 6.3 million people and kill 7,000 people each year (Ferrari et al., 2024; Gómez-Ochoa et al., 2022; Vucetich et al., 2025). *T. cruzi* is transmitted primarily via the triatomine vector, though blood transfusions, oral infection, and congenital transmission are appreciable sources of new cases. Chagas disease is a phenotypically heterogeneous disease; 30% of infected patients will develop potentially fatal cardiomyopathy, while others may develop megacolon or megaesophagus, and most will remain asymptomatically infected (Bern, 2015).

Accompanying this phenotypic diversity is a substantial amount of genomic variability. Parasites lineages are grouped into discrete lineages based on their nuclear genome sequences, referred to as discrete typing units (DTUs; TcI-TcVI), which segregate into clades that define the genomic ancestry of each strain (Freitas et al., 2006; Zingales et al., 2012). TcI and TcII are the most genetically distant from each other, while TcV and TcVI are both hybrids of TcII and TcIII. Substantial work has been done defining parasite diversity via simple genetic typing; however, relying on only parasite DTU to describe *T. cruzi* genomic variability potentially obscures important genomic variation that may contribute to clinical presentation. For example, although type TcV strains are especially common in areas with high clinical burden of Chagas diseases such as Bolivia, and are implicated in congenital transmission, there is substantial evidence of biologically relevant variability within this DTU (Callejas-Hernández et al., 2018; Hakim et al., 2023; Minning et al., 2011). Despite this, little work has been done characterizing the range of genomic diversity within TcV strains.

The extensive genomic variability even within parasite lineages is at least partially the result of *T. cruzi’s* remarkable genomic plasticity. Copy number variability has been observed in field isolates from vectors, and serial passage in culture has experimentally demonstrated the parasites’ capacity for copy number plasticity (Matos et al., 2022; Reis-Cunha et al., 2018; Sanchez et al., 2023; Vargas et al., 2004). *T. cruzi* genomes have core components that are largely syntenic across trypanosomatids, but these regions are disrupted by substantial regions that encode for large multigene families (Berná et al., 2018). These multigene families (MGFs) include mucins, mucin associated proteins (MASPs) trans-sialidases, discrete gene family-1 (DGF1), glycoprotien-63 (GP63) and retrotransposon hotspot proteins (RHSes), which are highly immunogenic and largely surface-exposed, and thus likely to be important for pathogenesis (Berná et al., 2024; Buscaglia et al., 2006; Durante et al., 2017; Fonseca et al., 2019; Risso et al., 2004; Teixeira et al., 2015)

Although it is clear that *T. cruzi* genomes are highly variable and diverse even within individual parasite lineages, these complexities have only been studied in strains adapted to laboratory culture. This is largely due to the challenges associated with generating sufficient genetic material required for whole genome sequencing of clinically isolated strains. Parasites are generally undetectable in the blood during the most common chronic stages of Chagas disease, and, even when parasites are detectable, isolating parasites from clinical specimens is a challenging and time intensive process. The few studies of clinically isolated specimens have demonstrated that they are phenotypically distinct from lab adapted strains (Engel et al., 1982; Faral-Tello et al., 2023; Veloso et al., 2005). The possible genomic basis for these phenotypic changes remains unclear.

In this study, we investigated the link between the genetic variability of *T. cruzi* and its phenotypic heterogeneity. To identify genetic features that could be important for clinical infection, we isolated and sequenced *T. cruzi* parasites from clinical cases of Chagas disease from three different clinical studies, representing different infection contexts. The participants included people with immune compromise due to HIV, pregnant people with Chagas disease sampled when they presented for delivery, and chronically infected participants in a Chagas cardiomyopathy study, representing clinical contexts with a range of age and immunosuppression. Our analysis revealed regions of variable copy number that may suggest parasite adaptations to the host environment and a repertoire of multigene family members in clinically isolated strains that is absent from lab adapted strains of the same genetic type. These results point towards genomic features of the parasite that may be critical for clinical infection, while also highlighting the importance of studying parasite genomic variability in a clinical context.

## METHODS

### Study information and ethics considerations

Samples were collected from three ongoing clinical studies based in Santa Cruz, Bolivia. The studies were approved by Johns Hopkins School of Public Health under Institutional Review Boards (IRB) numbers IRB00004303, IRB00005598, and IRB00002644 for opportunistic infections during HIV infection, Cardiomyopathy, and Congenital Studies respectively, as well as by local review boards at Hospital Universitario Japonés, Asociación Benéfica Proyectos en Informatica (PRISMA), and Universidad Catolica Boliviana. A description of larger congenital cohort can be found in Kaplinski et al and Messenger et al, while description of the Cardiomyopathy study can be found in Okamoto et al and Sherbuk et al (Kaplinski et al., 2015; Messenger and Bern, 2018; Okamoto et al., 2014; Sherbuk et al., 2015).

### Sample collection and culture preparation

15 mL of venous blood was collected in EDTA tubes from *T. cruzi* PCR positive patients Samples were centrifuged at 1200g for 10 mins at 4 degrees C. All but 0.5 mL of plasma and packed red cells was removed from the supernatant, then 8mL of supplemented RPMI-1640 was added to the packed red cells. Sample was spun again at 1200g for 10 mins, supernatant was discarded, and 2mL of packed red cells was distributed to three tubes filled with biphasic media containing blood agar (1.4% Blood agar, (Sigma 70133), 0.5% trypticase peptone (Sigma T9410), 0.6% purified agar and 0.6% NaCL) and overlay (0.9% NaCL and 150ug/ml gentamicin (Sigma G1522)). Biphasic media cultures were incubated at 24-28 degrees C, and cultures were examined for flagellates at 30, 60, 90, 120, 150, and 180 days. Once parasite growth was observed, cultures were transferred to Liver Infusion Triptose medium. After approximately 10 days of cultivation and once adequate proliferation was achieved, three washes with PBS were performed by centrifugation at 2000 g for 15 minutes. The resulting final pellet was resuspended in 10% DMSO in inactivated FBS and was stored at –80 °C until further processing.

### Sequencing and assembly

Samples were expanded and cryopreserved as previously, but all parasite cultures were non-viable upon thawing (Messenger et al., 2015). DNA was extracted using the QIAamp DNA blood kit - genomic DNA extraction (Qiagen, 51104), and Illumina libraries were prepared using the NEB Nextera XT DNA Library Preparation Kit kit (NEB FC-131-1096). Libraries were sequenced on an Illumina NovaSeq X at the Johns Hopkins Genetic Resources Core Facility (GRCF). Sequencing adapters were trimmed using bbduk (“BBMap,” 2025). Raw reads were assembled using Spades with kmer lengths of k = 21,33,55,77,99, and 127. To eliminate contaminating DNA from other organisms, a search for known *T. cruzi* sequences using BLASTN to query all assembled contigs against a concatenated library of the following high quality reference *T. cruzi* genomes representing different parasite genetic types: CL Brener Esmeraldo-like, CL Brener Non-Esmeraldo-like, G, BrazilA4, JRcl4, TCC, Tulacl2, Tulahuen, SlyvioX10, YC6, Bug2148, 231, Berenice, Dm28c and CL(Baptista et al., 2018; Berná et al., n.d.; Bradwell et al., 2018; Callejas-Hernández et al., 2018a; El-Sayed et al., 2005; Franzén et al., 2011; Hakim et al., 2024; Lewis et al., 2009; Wang et al., 2021). Raw reads were mapped to the pre and post-cleaned assemblies using bowtie2 (Langmead and Salzberg, 2012). Removal of contaminants was visualized in 2D histogram plots of GC content vs median contig coverages (**Supplemental Figure 1**). Raw fastq files were cleaned of contaminating sequencing reads using SAMtools (Danecek et al., 2021). The cleaned assemblies and fastqs were used in all further analyses. In addition to the clinical specimens, the following lab adapted strains were sequenced in house and processed in the same way to improve comparisons: Dm28cCas9, Tulahuen, and Colombiana.

### Bioinformatic analysis

Heterozygosity was assessed in clinical isolates using jellyfish to generate kmer histograms, with a kmer length of 21, then Genomescope (Marçais and Kingsford, 2011; Ranallo-Benavidez et al., 2020). Assembly statistics were assessed using Assembly-stats, BUSCO scores were calculated using Complasm (Huang and Li, 2023). All assembly statistics and sequencing statistics are reported in **Supplemental Table 1**.

FreeBayes was used to detect single nucleotide polymorphisms (SNPs) in clinical isolates against the CL Brener Esmeraldo-like reference genome using all default parameters, except for the parameters -F 0.01 -C 1 --pooled-continuous (Garrison and Marth, 2012). Allele frequency histograms were generated using these SNPs to assess complex infection, excluding sites with coverage less than 10.

Genetic distances between genome assemblies and known strains were calculated and visualized with Sourmash (Brown and Irber, 2016). Additional genetic distance among hybrid clinical isolates was assessed using the SNP frequency of clinical isolates against the TCC reference genome using InStrain, again excluding sites with coverage less than 10 (Olm et al., 2021). InStrain SNP data was also generated compared to the Cl Brener Esmerldo-like reference genome and was used to estimate levels of heterozygosity in 5kb genomic windows, excluding sites with more than 2 alleles and coverage less than 10.

Copy number variation was assessed as previously described (Matos et al., 2022). Raw reads of each sample were mapped against the CL Brener Esmerldo-like reference, and Control-FREEC was used to estimate copy number of 5 kilobase windows across the reference genomes (Boeva et al., 2012). In order to rank genomic windows in order of copy number variability, we calculated, for each window, the median absolute deviation of normalized read depth coverage at the same site in every sample. We used a Fisher’s test to assess the significance of multigene family (MGF) enrichment in the top percentile of variability.

Multigene family members were annotated from clinical isolate assemblies, short read assembled lab strains, and previously generated reference genomes, using the same methodology previously described (Hakim et al., 2024; Wang et al., 2021). The identified genes were clustered using CD hit at 98% sequence identity using a word size of 9. After clustering, genes within a cluster shorter than 500 base pairs were removed, to increase specificity and to avoid erroneous clustering of fragmented assemblies. A schematic representation of this approach is in **Supplemental Figure 2**.

BLASTN was used to search for MGFs assembled from clinical isolates that were putatively lost from lab adapted strains. Using the representative sequence from the gene cluster as the subject queried against TcII and hybrid reference strains, all MGFs members that matched an MGR sequence with at least 98% sequence similarity and 50% query coverage to were considered present in the lab strain.

Statistical analysis was done in R v.4.2.2. All code to run analysis and generate figures are available at https://github.com/mugnierlab/Hakim_2025. Raw sequencing data is available at SRA BioProject PRJNA1271783.

## RESULTS

### Clinical isolates from different infection contexts in Bolivia display underlying genetic variability

*Trypanosoma cruzi* was cultured from three clinical studies in Bolivia (**Supplemental Table 1**). Hemoculture was attempted from 148 patients with confirmed positive *T. cruzi* serology yielding a total of 18 positive hemocultures, resulting in a disaggregated success rate of 5.61% (5/89) from peripartum mothers, 19.57% (9/46) from chronically infected patients, and 33.34% (4/12) from participants with end stage AIDS. Of these samples, 15 isolates across the three studies had sufficient material and were sequenced using Illumina short read sequencing.

*T. cruzi* infection can be multiclonal, especially in immunosuppressed individuals, such as pregnant people and people living with HIV (Bowman et al., 2021; Hakim et al., 2023; Talavera-López et al., 2021). To assess the clonality of each sample, we mapped the raw reads of each isolate against the CL Brener Esmeraldo-like reference genome. We observed allele frequencies that centered around 0, 1 and 0.5 in all sequenced samples, indicating the presence of a single diploid organism (**Supplemental Figure 3).** Based on these results, we conclude that, although the original patients may have had polyclonal infections, these clinical samples each represent a single parasite clone.

To identify parasite lineages, we used SourMash to plot the Jaccard distance of kmers between the assembled clinical isolates and reference genomes of known DTU designation (**Figure 1A**). Of the 15 isolates, 13 clustered most closely to the hybrid TcV reference genome, while the remaining two clustered with type TcII reference genomes. Moreover, other work in the same Bolivian population has shown a predominance of type TcV strains (Bern et al., 2009). In the absence of a high-quality genome for *T. cruzi* DTU V, we mapped the raw reads from the hybrid strains to the best available reference genome for hybrid lineages (TcVI TCC genome) to assess additional genetic variation. We called SNPs to generate a PCA plot of allele frequencies to estimate genetic distance between parasites of the same genetic type **(Figure 1B)**.

**Figure 1:**
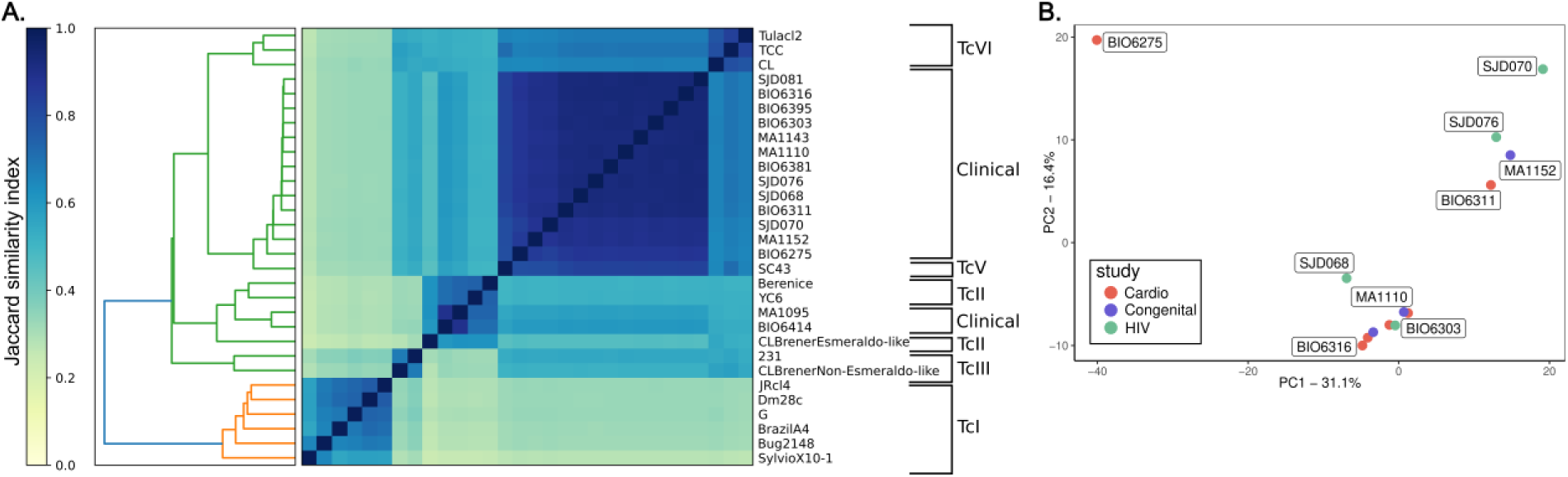
15 clinical isolates from different infection contexts in Bolivia display underlying genetic variability. A) Jaccard similarity matrix of kmers across assembled genomes. Yellow is most dissimilar and blue is the most similar in kmer overlap. Hierarchical clustering of the data from the Jaccard matrix is shown in the dendrogram. B) PCA of allele distances among genes in hybrid genomes.

In addition to varying at the SNP level, *T. cruzi* and other trypanosomatids are also known to undergo gene copy number variation (Bussotti et al., 2018; Laffitte et al., 2016; Matos et al., 2022). To evaluate potential copy number variations associated with clinical infection, we measured the level of copy number variation across 5kb windows in our samples using Control-FREEC. We found regions of copy number variation across all contigs and all samples (**Figure 2A**). Some genomic regions, such as those that map to Chr31, have increased copy in all clinical samples, which has been previously observed in field isolates and may represent a universal genomic feature of field isolated *T. cruzi* (Reis-Cunha and Bartholomeu, 2019; Sanchez et al., 2023). Of particular interest are sites that have variable copy number in multiple clinical isolates, because they may represent recent or specific adaptations to the clinical infection context; for example, some of the genomic information encoded in chromosome 9 with elevated copy number in the MA110 genome shows decreased copy number in the MA1143 genome. In other work, sites with variable copy number are often enriched for multi-copy genes including *T. cruzi’s* highly antigenic multigene family members such as retrotransposon hotspot proteins and trans-sialidases (Matos et al., 2022; Reis-Cunha et al., 2018; Talavera-López et al., 2021). To formally assess whether genomic regions that are consistently variable in copy number are enriched for MGFs, we used a Fisher’s test to assess MGF enrichment in the most variable genomic windows. We ranking genomic windows based on copy number variance, we found that the most variable windows, those in the top 5^th^ percentile, are significantly more likely to be enriched with MGF members, compared to the background distribution of the whole genome (OR: 3.29 for presence of MGF genes in high CNV variance windows, CI 3.26-3.34). This observation is consistent with other analyses; in long read genome assemblies, MGFs have variable repertoire sizes in different *T. cruzi* strains (Callejas-Hernández et al., 2018b; Matos et al., 2022; Reis-Cunha et al., 2022; Wang et al., 2021).

**Figure 2:**
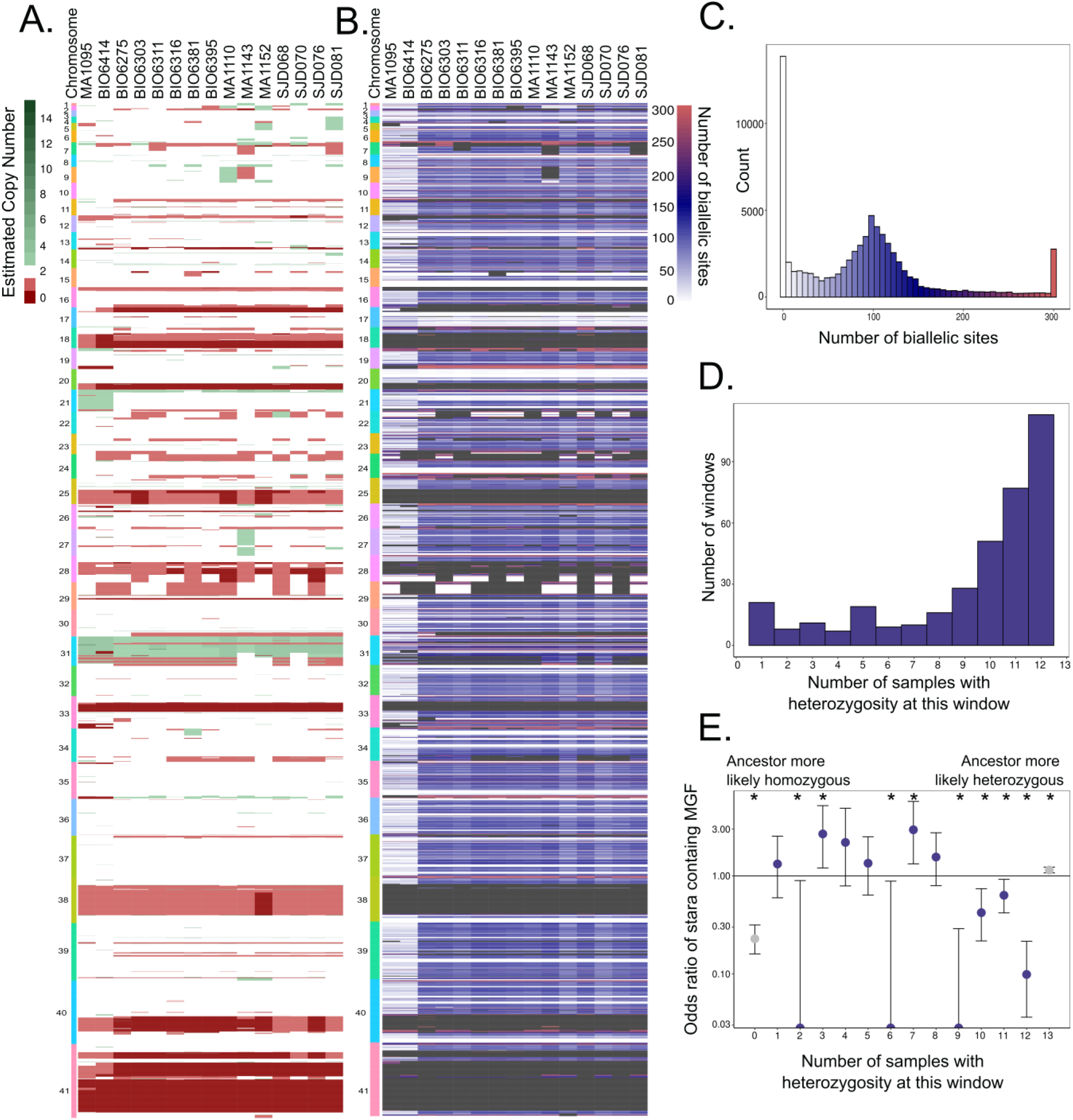
Variable copy number across genomic regions and patterns of homozygosity suggests genomic diversification. **A**) Estimated copy number variation for 5 kb windows in clinically isolated parasite genomes. Each row is a 5kb window colored by loss (red colors) or gain (green) of genomic content at that site. Copy number gains are censored at 15. **B)** Sum of sites in each genome with exactly two alleles in each 5kb window. White indicates regions with very few biallelic sites, indicating homozygosity, while blue then red indicates an increase of biallelic sites. Windows are censored at biallelic sites = 300. Grey boxes are masked regions that have less than two estimated copies, which would by definition have one copy of the allele. **C)** Histogram showing the distribution windows with a given number of biallelic sites. The heterozygous peak is largely at 100 biallelic sites per 5kb, or 2%, and therefore a minimum threshold of 1% of biallelic sites is set for calling a window heterozygous. The colors correspond to **figure 2B**. **D)** Heterozygosity concordance among TcV samples. For each window, the number of samples where that window is largely heterozygous (greater than 1% of sites in the window are biallelic) was calculated. The number of windows that are heterozygous in one out of 13 total samples TcV samples, up to 12 out of 13 TcV samples is shown. The majority of windows had perfectly concordant heterozygosity (n = 1035) or concordant homozygosity (n = 2985). The sites without perfect concordance in heterozygosity in all TcV samples are marked as possible gene conversion events, in blue. **E.** The odds ratio of a window in each stratum defined in **figure 2D** to have MGF members. Windows with 2, 6 and 9 samples sharing heterozygosity had zero observed MGFs and thus the CI reaches 0. Asterisks indicate ORs that are statistically significant after correction via Benjamini-Hochberg Procedure. ORs and CIs are available in Supplemental Table 3.

Variance in copy number for specific genomic regions raises the question of overall genomic content, both in terms of haploid genome length and proportion of the genome that contains heterozygous alleles. To generate a reference free estimation of genome length and heterozygosity, we generated kmers with jellyfish and analyzed the kmer histograms with GenomeScope. Despite the copy number variation observed in **figure 2**, the total genomic material remains similar across all isolates (37Mb – 44Mb), around the expected range for *T. cruzi* (40 Mb; (Souza et al., 2011)). As expected, the two non-hybrid TcII samples, MA1095 and BIO6414, have lower heterozygosity (0.98% and 1.04%), while the hybrid TcV samples have high heterozygosity (2.35% - 2.59%) **(Supplemental Table 1)**.

We investigated the heterozygosity of the clinically isolated genomes more closely by counting the number of bases in the genome that contain 2 alleles within a 5kb window (Figure 2B**,2C**). Windows with more biallelic sites have higher heterozygosity, and those with more monoallelic sites have higher homozygosity. We find regions of both shared and variable heterozygosity between all genomes, which could represent adaptation, either by diversity generation or elimination, in these parasites.

To more closely evaluate potential adaptation in these isolates, we analyzed genetic windows in the TcV clinically isolated genomes that vary in their level of heterozygosity (**Figure 2D**). Given the shared lineage of these parasites, this discordance of heterozygosity must have arisen after these strains diverged from a common ancestor. Discordant windows may have either started out homozygous, later becoming heterozygous by random mutation in only one allele or by homologous recombination with another non-allelic locus to generate additional diversity, or alternatively, started out heterozygous and became homozygous by allelic homozygous recombination, using the alternate allele as a template. Using the number of samples with shared heterozygosity as a proxy for likelihood that a common ancestor originally had heterozygosity in that window, we see that regions that are heterozygous in most samples and thus were likely heterozygous before conversion to homozygosity in a subset of samples, are highly depleted of MGFs compared to the background distribution of MGF genes in the genome as a whole. (windows where 9, 10, 11 and 12 out of 13 samples are heterozygous at that window, OR = 0 (0-0.20), 0.37(0.29-0.62), 0.50 (0.33-0.71), 0.21 (0.12-0.33)). In contrast, we do not observe consistent depletion of MGFs in regions where the majority of sites are homozygous and likely ancestrally homozygous. These observations suggest that recombination is being used both as a mechanism to increase diversity in MGF-containing genomic regions via copy number variation and to eliminate or limit diversity via homozygosity-generating gene conversion in important core genomic features.

### Multigene family repertoires in *T. cruzi* clinical isolates are distinct from the repertoires of strains adapted to lab culture

The expansion of MGFs in the *T. cruzi* genome is a major source of complexity and has rarely been investigated in clinically isolated strains. In high quality genome assemblies of reference strains adapted to lab culture over long periods of time, the repertoires of these genes are dissimilar (Wang et al., 2021). As these genes are important for infection and likely pathogenesis, it is possible that the repertoire maintained in lab-adapted *T. cruzi* strains may be distinct from those in clinically isolated strains. In order to assess MGF repertoire membership in the assembled *T. cruzi* genomes, and to determine if there were any differences in repertoire between clinically isolated and lab adapted strains, we annotated the clinical isolate assemblies for MGFs using a BLASTN approach against a library of annotated MGFs from the Cl Brener genome, as previously described (Hakim et al., 2024; Wang et al., 2021). We repeated this annotation strategy for the lab adapted strains, using previously published long and short read assemblies, as well as in-house generated short read assemblies, to generate a comparable dataset. To measure the diversity of MGFs in fragmented short read assemblies, we clustered all the genes within a multigene family from all samples together using CD-HIT at 98% similarity. This approach should consolidate most small assembly and PCR errors, allowing the identification of related or identical MGF clusters that were present in multiple samples. **(Supplemental Fig 2)** This approach yielded gene clusters within each multigene family that were present in both clinical and lab adapted strains.

In order to assess the repertoire of MGF gene clusters across samples, we visualized the presence or absence of a member of a gene cluster within each sample with a Jaccard similarity matrix and plotted the results on a multidimensionality scaling plot (**Fig 3A**). The samples clustered based on DTU, as expected, but did not cluster based on the specific clinical context. Instead, we saw a striking difference between lab-adapted and clinically isolated strains. To better understand how the MGF repertoire differs between lab-adapted and clinically isolated strains, we next identified gene clusters present in all clinical isolates. We were surprised to find that there was a subset of gene clusters found in every clinical isolate that was also absent from all the queried lab adapted strains, regardless of DTU (**Figure 3B**). To more rigorously determine whether these MGFs were truly unique to clinical isolates, we used BLASTN to search for the representative sequence from each gene cluster (the longest sequence within each cluster) within the lab adapted strain’s genomes (**Supplemental Figure fig 3A and B**). After this filtering out MGFs present in any lab strain (>98% sequence similarity and 50% query coverage), we identified between 15 and 47 genes per multigene family that were unique to clinical isolates and absent from all laboratory strains (DGF1= 15, GP63 = 20, mucin = 28, RHS = 16, Tran-sialidase = 40, MASP = 47). To assess the statistical significance of this observation, and to confirm it was not an artifact of the number of gene clusters we were able to assemble and annotate in each sample type, we permuted the gene clusters found in clinical isolates and the genetically related type TcII, TcV and TcVI lab adapted reference strains to estimate the number of gene clusters that we would expect to be present in all clinical isolates and no lab strains by chance **(Figure 3C)**. Across all gene families, the observed frequency of unique gene clusters for each MGF was highly significant as calculated by z score (zscore of DGF1 = 12.17, GP63 = 199.9, MASP =333.38, mucin = 279.9, RHS = 28.7, TS = 74.93, all p values by the z statistic 0.00001). Thus, the high number of genes that were absent from lab strains but present in all clinical isolates is unlikely to have occurred by chance. Our current understanding of *T. cruzi* genetic ancestry would suggest that these gene clusters, exclusive to a set of clinical isolates that includes both TcII and hybrid strains, are likely to have been present in a common ancestor of both lineages (Zingales et al., 2012). They were therefore likely later lost from the lab-adapted strains during adaptation to culture and may represent gene clusters that are important for clinical infection but dispensable for growth in culture. We did not find any shared sequence motifs in the translated representative sequences of the clinically conserved gene clusters.

**Figure 3:**
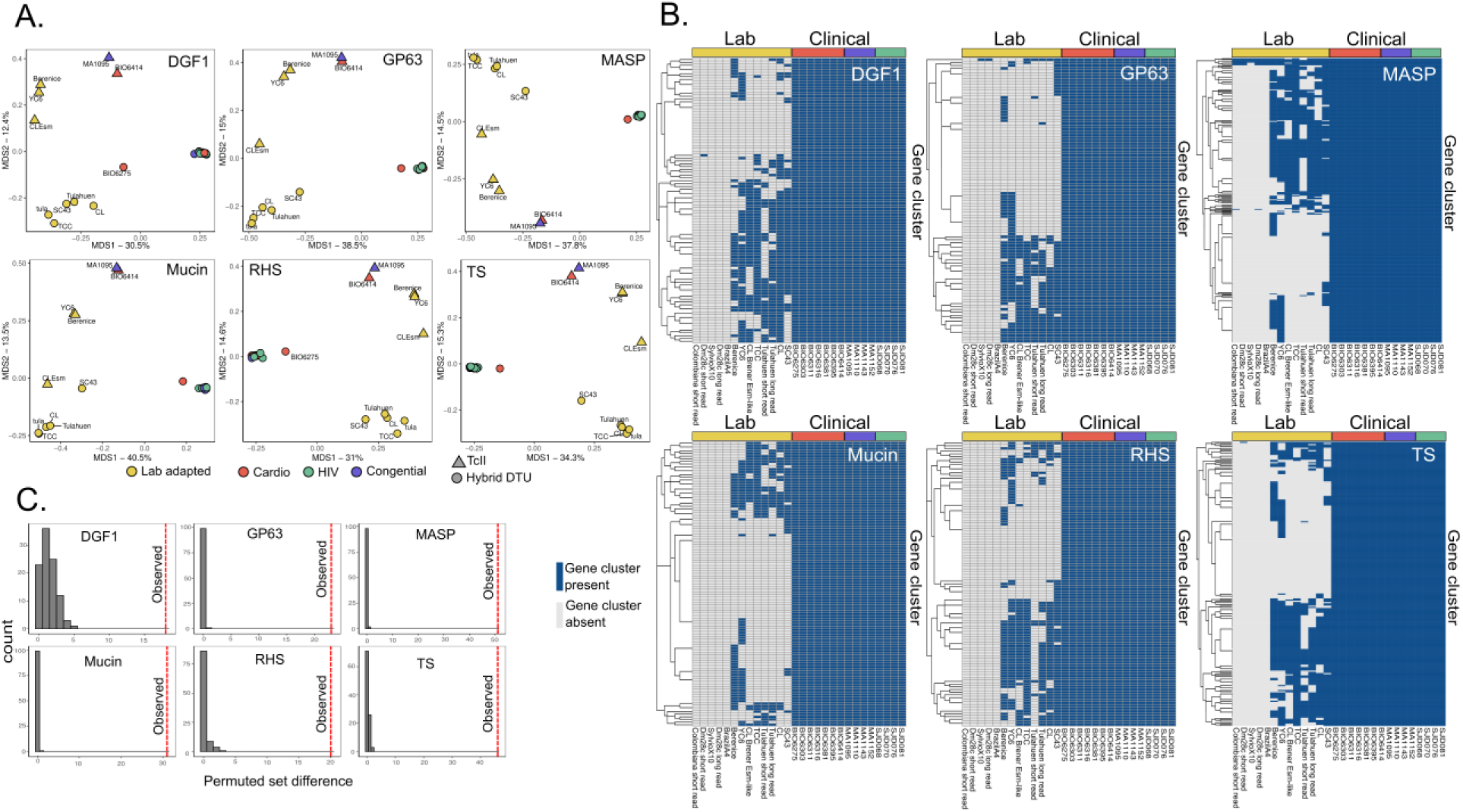
Multigene family repertoires in clinically isolated strains are distinct from the repertoires of strains adapted to lab culture. A) Multidimensional scaling plot of the Jaccard distance between samples’ MGF membership visualizing the presence or absence of a member of a gene cluster within each sample. Color of points indicates study, and shape indicates parasite genetic type. B) MGF clusters that are found in all clinical isolates, regardless of DTU. Columns are strains, and those indicated with yellow bars are lab adapted genomes. Blue cells are gene clusters that were present in the genome, grey cells were absent. C) Histogram of permuted set difference, establishing the null distribution for set difference between the clinical isolates and lab adapted TcII, TcV, and TcVI strains. The dotted red line represents the observed set difference between the sets of MGF clusters found in all clinical isolates and the set of MGF clusters found in none of the lab adapted TcII, TcV or TcVI strains.

We excluded TcI reference genes from MGF repertoire comparisons because our study did not have any clinically isolated specimens that were type TcI (**Supplemental Figure 4A)**. Moreover, because TcI parasites are generally more genetically diverse, and may share a more ancient common ancestor compared to TcII and TcV parasites, we may expect their MGF repertoires to be more diverged. To assess if these findings are generalizable to TcI parasites, we repeated the analysis using previously published whole genome sequencing data from parasites isolated from Ecuadorian vectors and from two Colombian patients, one with Chagasic megacolon and one with cardiomyopathy (Schwabl et al., 2019; Talavera-López et al., 2021). We did not observe the same separation between clinical and lab adapted strains (**Supplemental Figure 4B)**. We also found no overlap between the MGF members shared in our Bolivian clinical isolates and the MGF repertoires of other TcI *T. cruzi* field isolates from vectors, suggesting these genetic types are too separated in evolutionary time to share MGF gene clusters.

To further interrogate the overlap of MGF membership within each DTU, and to observe if TcI MGF repertoires were more diverged than the repertoires of hybrid strains, we counted the number of parasite genomes a MGF gene cluster was found in. We observe that MGF gene clusters found in TcI lab strains are more likely to occur in one or two lab strains, while gene clusters found in hybrid DTUs are more likely to occur in 2 or more, suggesting increased MGF divergence in TcI parasites. This increased divergence in MGF repertoires may explain the lack of clustering we see among TcI strains (supplemental figure 4B). Despite this divergence, we observe conservation of the MGF repertoires among field isolated strains in both hybrid and TcI DTUs, with recent field isolates showing greater repertoire overlap than those adapted to culture **(Figure 4)**. While we cannot compare the size of MGF repertoires in samples, this suggests some preference for the maintenance of a larger MGF repertoire in circulating parasite strains.

**Figure 4:**
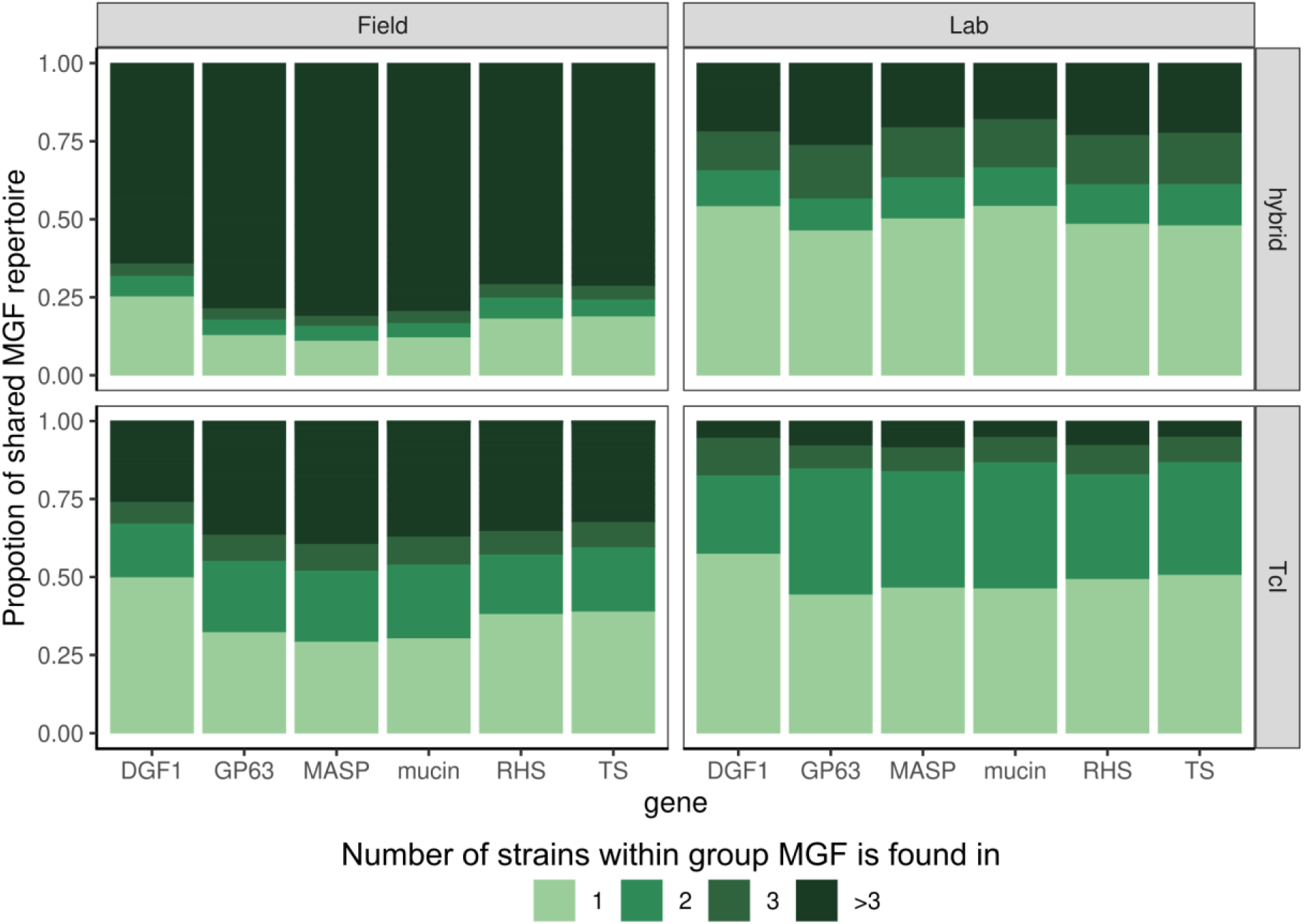
MGF repertoires are more conserved in recent field isolates compared to laboratory adapted strains, regardless of DTU. Proportion of the MGF repertoire that is shared across DTU and isolation recency. For each MGF, the number of field isolated TcI, lab adapted TcI, field isolated hybrid, or lab adapted hybrid strains that each gene cluster is found in is graphed in a proportional bar graph.

## DISCUSSION

In this study, we isolated and sequenced *T. cruzi* genomes from a diverse population of patients living with Chagas disease. This study provides the first whole genome sequencing data for clinically isolated hybrid strains, demonstrating regions of variable copy number across clinical isolates, potential sites of gene conversion, and MGF repertoires that are distinct from strains that have been adapted to laboratory culture. Finally, we observe a set of MGF member gene clusters that are conserved across Bolivian clinical isolates, regardless of parasite genetic ancestry, that appear to have been lost in parasite strains adapted to laboratory culture; similarly, we observe a general decrease in MGF repertoire conservation among lab adapted strains compared to recently isolated parasites that is similarly independent of DTU. By comparing the genomes of recent clinical isolates to those of lab adapted strains, we have identified features of the parasite genome that might be critical for clinical infection, highlighting the value of understanding *T. cruzi* genomes in the field.

Our unbiased whole genome approach allows a holistic investigation of genome diversification for each isolate. *In vitro* studies have demonstrated the genomic plasticity of *T. cruzi*, and previous work has demonstrated genome copy number variation and the possibility of meiotic recombination in recent vector isolates (Berry et al., 2019; Schwabl et al., 2019; Talavera-López et al., 2021). Here, we find similar plasticity in parasites isolated from human infections. These variable regions point towards genes or sets of genes that may be important for adaptation to the host environment. We find genomic regions gaining homozygosity are depleted of antigenic genes, whereas regions with copy number changes are enriched for them. Thus, we observe a compartmentalization between the maintenance and the diversification of genomic content, with diversifying patterns of genome evolution enriched in regions encoding for MGF members, which are involved in immune evasion, and maintenance patterns enriched in regions encoding for core genomic features.

Recombination following double stranded DNA breaks is a possible mechanism for both the generation and elimination of diversity that we observe. Because *T. cruzi* lacks non-homologous end joining, DNA double strand break repair in trypanosomes occurs via either homologous recombination or microhomology mediated repair (da Silva, 2021). In higher order eukaryotes, the template used for homologous recombination is most often the alternate allele on the homologous chromosome, leading to gene conversion that eliminates allelic diversity by removing heterozygous alleles from an individual and a population (Richardson et al., 1998). We observe patterns consistent with this mechanism in regions depleted of MGF members. Increased copy number regions, in contrast, are likely to arise from non-allelic recombination, where unequal crossing over occurs. These regions of unequal crossing over may be templated by highly repetitive transposable elements. We and others have previously observed that MGF members are closer in linear genome space to transposable elements than other gene groups (Hakim et al., 2024; Macías et al., 2018; Ribeiro et al., 2019; Talavera-López et al., 2021; Vázquez et al., 2000; Wang et al., 2021). Antigenic gene copy number may increase over time by gene drift, increasing the potential genomic repertoire for these diverse genes. Because *T. cruzi* is thought to lack transcriptional control, gene dosage could also be a genomic adaptation resulting in phenotypic changes (Matos et al., 2022). Regions with variable copy number in clinical isolates, including those containing non-MGF genes, may be specifically important to clinical infection. The compartmentalization of diversity-generating and diversity-eliminating recombination may be driven by the increased rate of non-allelic recombination near MGFs given the abundance of potential recombination templates, either from other MGF members or from transposable elements proximal to MGFs, or could be a result of negative selection against deleterious diversity in core genomic regions. Both outcomes are likely to contribute to this observation.

In this study, we were originally interested in observing differences in parasite genomes related to specific infection contexts. The patients in the study represent a gradient of immunosuppression, with varying levels of parasitemia, age, and severity of disease. Even though these sequenced parasites have likely been circulating within each individual for several years and in some cases decades, providing ample time for the parasites to accumulate common mutations related to the specific infection context, we were unable to observe differences in nucleotide diversity, MGF repertoire, or copy number variation specific to clinical infection context. It is important to note, however, that the samples from this study all represent chronic infections. Although pregnancy and acute reactivation due to HIV are associated with immunocompromise, this occurs over a relatively short timeframe during which unique genomic changes are unlikely to arise. Our inability to define genotypes associated with specific clinical contexts may also be due to the very small sample size, with as few as four isolates per group, which was underpowered to detect these differences in a whole genome approach. However, regions that we identified as genomically plastic across our isolates may be the focus of future targeted analyses that do not require the intensive strain isolation procedure employed here.

Although parasite genomes did not cluster genomically based on infection context, we did observe an underlying genetic structure unrelated to infection context. While it is unclear what features underlie this genetic structure, they are likely related to other epidemiological factors, most notably the geography of where the patients were initially infected. Previous work has shown that parasites isolated from vectors genomically cluster based on geographic distribution, even in within a single country (Berry et al., 2019; Schwabl et al., 2019). As the time of initial infection for patients in this study, likely many years prior to sampling, is unknown, it is nearly impossible to account for geographic distribution in our analyses. A larger study, or one specifically designed to control for geographic variability, may better reveal genomic features associated with specific clinical phenotypes.

Although we saw no changes associated with specific clinical phenotypes, we saw remarkable differences between clinical and lab-adapted strains. Our results suggest that MGF repertoires diverge during adaptation to lab conditions, with decreased overall conservation of MGF repertoires among lab-adapted parasite strains and a subset of MGF genes that are consistently lost in these strains but maintained in Bolivian clinical isolates. Notably, we did not find any of these conserved MGF gene clusters in previously published TcI data, likely due to the genetic distance between TcI and TcII/hybrid *T. cruzi* strains. We were unable to accurately describe the repertoire size of MGF members for clinical strains to compare them to lab adapted strains, due to well characterized issues in assembling multicopy and repetitive regions using short reads (Callejas-Hernández et al., 2018). However, for the MGFs that we were able to assemble and annotate, the number of gene clusters that were found in all clinically isolated strains and absent from all TcII or hybrid lab strains was much higher than would be expected by chance. While we cannot account for unassembled MGFs, it seems there is either active pressure to maintain these specific genes in clinically infective strains, or to eliminate them from lab adapted ones. Notably, several groups have observed that *T. cruzi* strains develop decreased virulence after serial passage in culture, potentially suggesting a role for these MGFs not only in clinical infection, but also in parasite virulence (Alves et al., 1993; Brener and Chiari, 1965; Veloso et al., 2005). The loss of MGFs, as well as parasite virulence, after passage in culture may be explained by a lack of evolutionary pressure to maintain these antigenically variable genes, as parasites in culture are often passaged in the absence of exposure to the host immune system. The maintenance of a conserved antigenic gene repertoire among genetically related strains suggests that there may be common features in conserved MGFs that are essential for infection. While we did not find any shared sequence motifs and the genes did not cluster together phylogenetically, long read genome assemblies from additional samples may help elucidate the biological mechanisms driving this conservation.

Aside from potential mechanistic implications, this shared repertoire of immunogenic genes may provide a promising avenue for the development of new, targeted multi-antigen diagnostics, which could outperform current diagnostic modalities. It is increasingly appreciated that differences in parasite genetics may drive lower serological test sensitivities, a major issue in Chagas disease diagnosis, which is primarily performed using serological tests. MGF members encode for highly immunogenic proteins, and recombinant serological assays often use motifs from these genes as targets. Serological tests have the lowest sensitivity in Mexico and Central America, regions where TcI parasite strains are predominant (Truyens et al., 2021; Velásquez-Ortiz et al., 2022; Whitman et al., 2019). In line with this, we observed very little overlap between MGF repertoires of TcI field isolated parasites and TcII/TcV clinically isolated parasites, potentially explaining the failure of serological tests in TcI predominant regions.

However, while we observe differences in MGF repertoire in TcI versus hybrid DTUs, we also observe an increased repertoire overlap in different field isolated strains of the same DTU. A better understanding of the overlap of MGF repertoires within each DTU may uncover conserved immunogenic genes within that parasite group, enabling the discovery of new serologic targets and the development of more sensitive tests.

An unavoidable limitation of this work is that we were only able to sequence parasite strains that were able to grow in culture. It is therefore possible that there is additional diversity in clinical isolates that we could not capture, and that some genes especially critical to clinical infection, potentially impeding parasite survival in culture, could not be studied under these circumstances. Our recovery of only a single parasite clone for each sample is likely a result of the bottleneck associated with serial passage in culture, as previous work has demonstrated the multi-clonality of many clinical Chagas infections, especially immunocompromised people (Bowman et al., 2021; Oliveira et al., 2021). Additionally, the surviving clones from each participant may have genomically adapted to culture during the 50-100 days the parasite was kept in culture, though this length of time represents very few cell cycles and thus limited opportunity for genomic changes. It is also important to note that, though the samples were isolated from a variety of clinical contexts at larger tertiary hospitals, the patients were still all from Bolivia, likely infected in the Bolivian Chaco or the inter-Andean valleys of Cochabamba, whereas the range of *T. cruzi* spans across Latin America. Future studies could evaluate the conservation of these genes in a larger and more geographically diverse sample set.

In Santa Cruz, Bolivia, where these parasites were isolated and where the hybrid TcV strain is predominant, one in five people has Chagas disease (Bern et al., 2009). Despite its public health importance, *T. cruzi* was the last of the clinically important trypanosomatids to have a complete genome assembly, due to its genomic complexity. As technologies improve, the extent of genomic variability that exists within *T. cruzi* strains is becoming more apparent (Callejas-Hernández et al., 2018; Matos et al., 2022; Reis-Cunha et al., 2022; Sanchez et al., 2023; Wang et al., 2021). Despite this growing body of knowledge, little is known about specific virulence factors that may direct pathogenesis, or account for the range of phenotypic outcomes of Chagas disease. Hybrid parasite strains and parasites isolated from clinical contexts have been especially neglected. This work presents the first investigation into the genomes of hybrid *T. cruzi* clinical isolates, which are the predominant strains in the highly endemic southern cone of the Americas. The data presented here, therefore, represent a critical first step in demonstrating the ways that parasites isolated from the clinic are biologically distinct from those that have been adapted to culture. Understanding these differences will be essential to developing more sensitive diagnostic tools and identifying new drug targets for this important neglected tropical disease.

### Data Sharing

All cleaned genome sequencing data have been deposited to SRA BioProject PRJNA1271783. All code to generate analysis and figures are available at Github https://github.com/mugnierlab/Hakim_2025.

### Funding Sources

This study was supported by NIH Global Research Training Grant D43 TW007681, NIAD grant R01-AI087776, A Johns Hopkins University Discovery Award, the Biotechnology and Biological Sciences Research Council Doctoral Training Grant, The Dr. Gordon Smith Travelling Fellowship, The Chadwick Trust Travelling Scholarship, and Royal Society of Tropical Medicine and Hygiene small grant. J.M.C.H. is supported by T32AI007417 and a JHU Discovery Award. C.D. is supported by F31HL173973. E.M.M. is supported by D43TW001140.

---

**Supplemental Figure 1:**
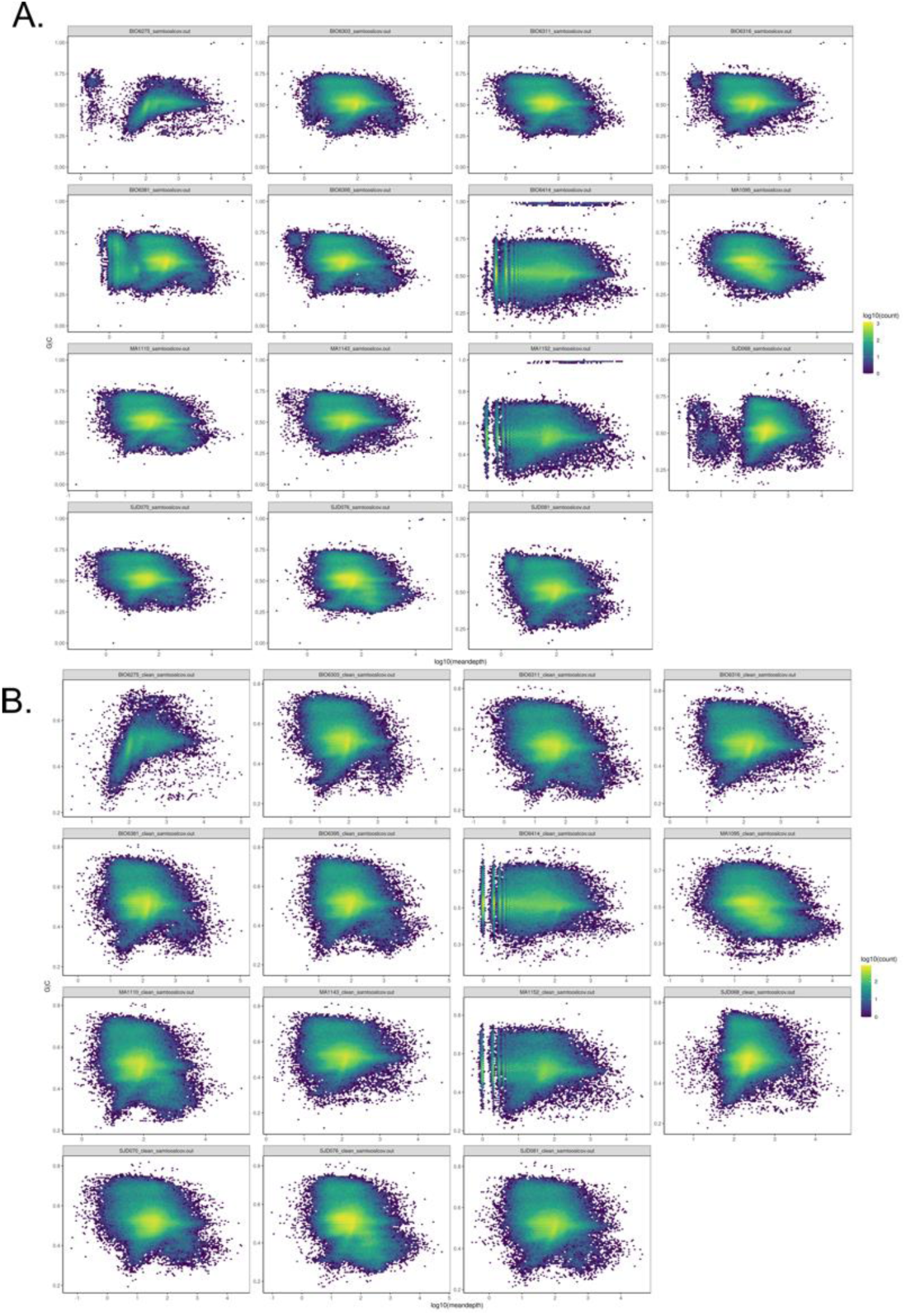
2D histograms of GC percent and contig coverage for clinical isolate assemblies before (A) and after (B) cleaning contaminating DNA and kDNA

**Supplemental Figure 2:**
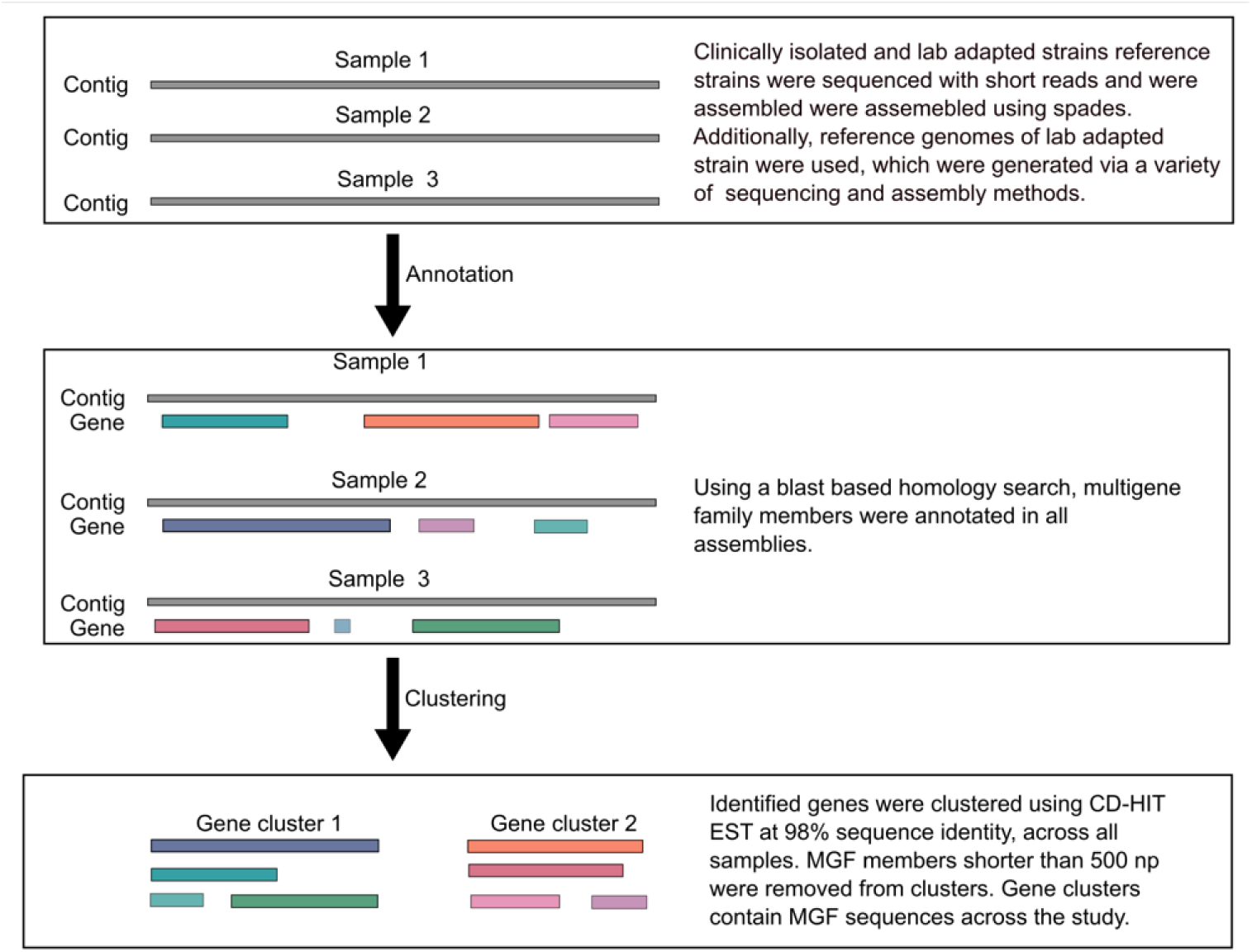
Schematic representation of gene clustering strategy for multigene family members

**Supplemental Figure 3:**
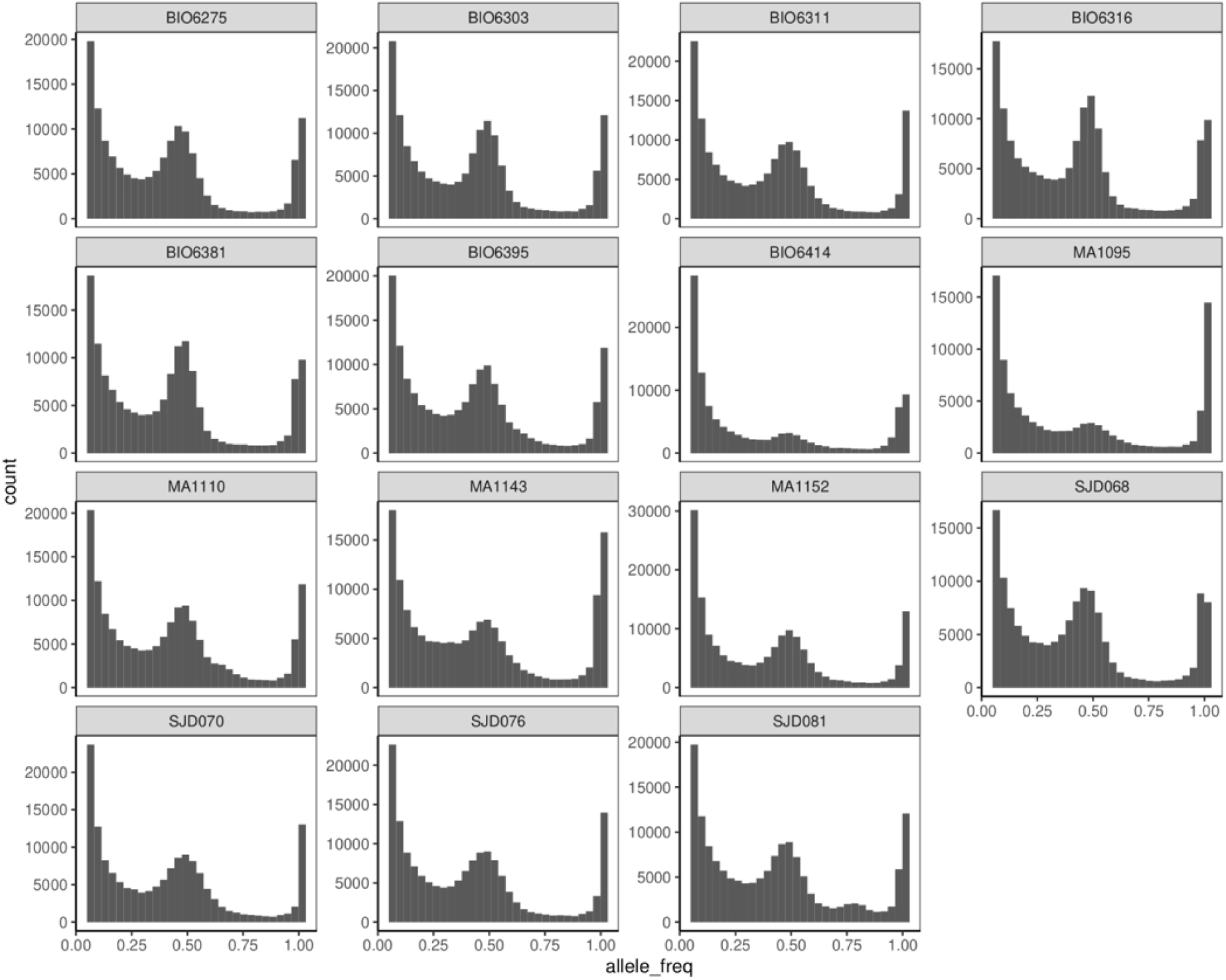
Distribution of allele frequency for clinically isolated strains against the CL Brener Esmeraldo-like reference genome. Isolates BIO6414 and MA1095 have smaller peaks at 0.5, due to their non-hybrid nature, and lower prevalence of heterozygous alleles.

**Supplemental Figure 4:**
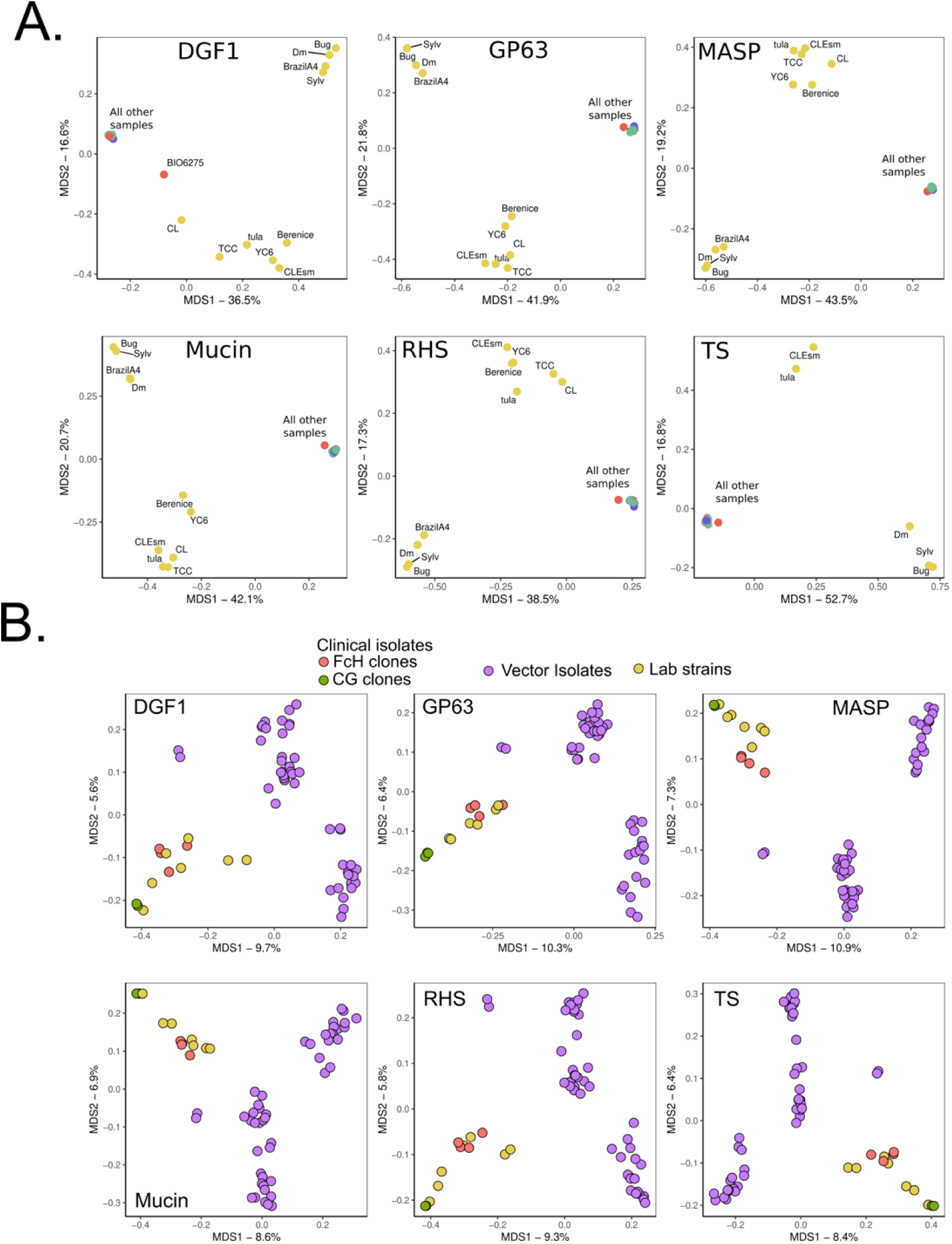
A) MGF repertoire separation including Tc1 lab reference strains. DTU separation is more pronounced but note that due to the lack of clinically isolated specimens in the current study some separation may be largely DTU driven. B) MGF repertoire separation of TcI parasites alone, including Colombian clinically isolated parasites from Talavera-Lopez et al (FcH and CG, red and green respectively) and vector isolates from Schwabl et al. (purple). FcH and CG are two individuals for which there were four and two parasite genomes cloned from the patient samples

**Supplemental Figure 3:**
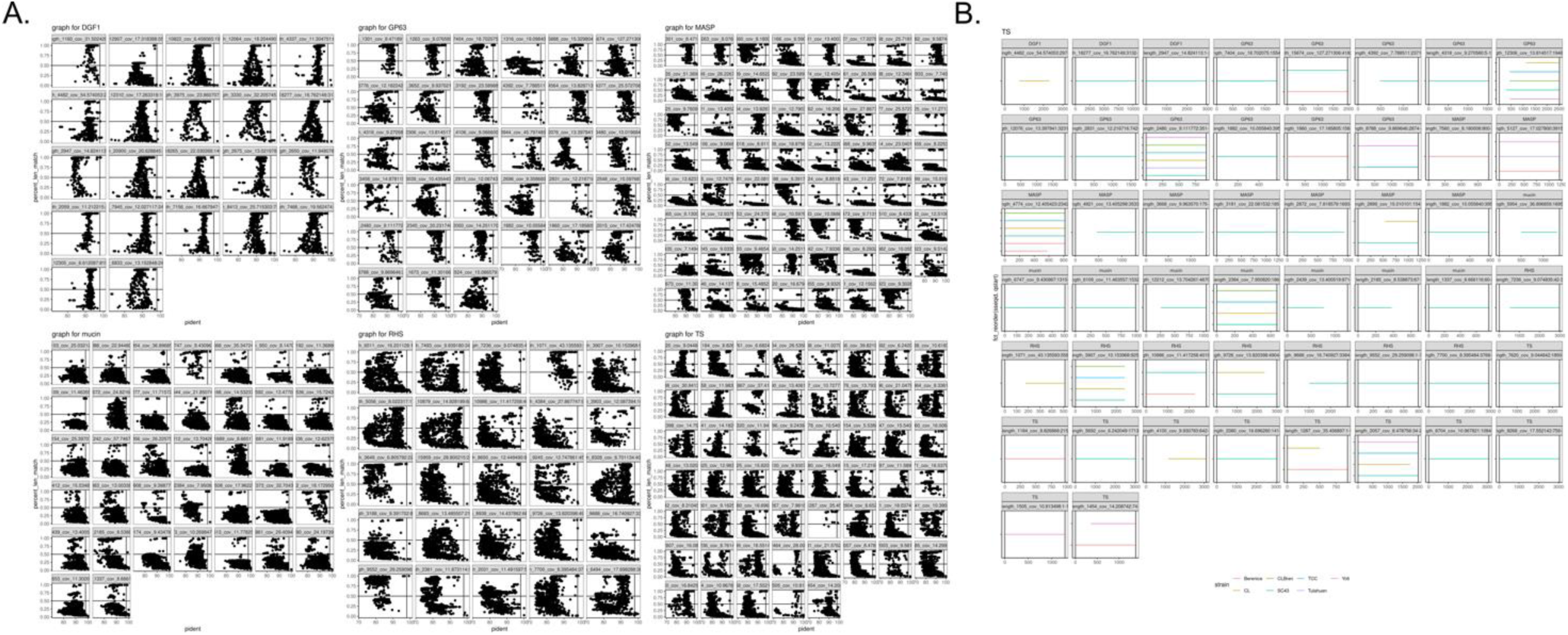
Confirming the loss of shared MGFs among lab adapted strains by BLASTN search of lost representative sequences within whole genomes of TcII, TcV and TcVI strains. **A)** Plot of all BLASTN hits in reference genomes for each lost MGF encoding gene, across all gene families. X axis represents the percent identity of the BLASTN hit, and Y axis represents the percent of the total MGF that is covered by the BLASTN homology found in the reference genome. The horizontal line is at 50% query coverage, and the vertical line is at 98% BLASTN sequence identity. **B)** Each MGF member that passes the above thresholds. Each gene is a facet, colored by the reference genome the gene was found in. X axis is the length of the gene, with black bars demarcating that start and end, and line segments of each reference genome representing the position where the hit occurred. These are the genes that were removed from the set of shared clinical MGF members that were lost in the lab adapted strains.

**Supplemental Table 1:**
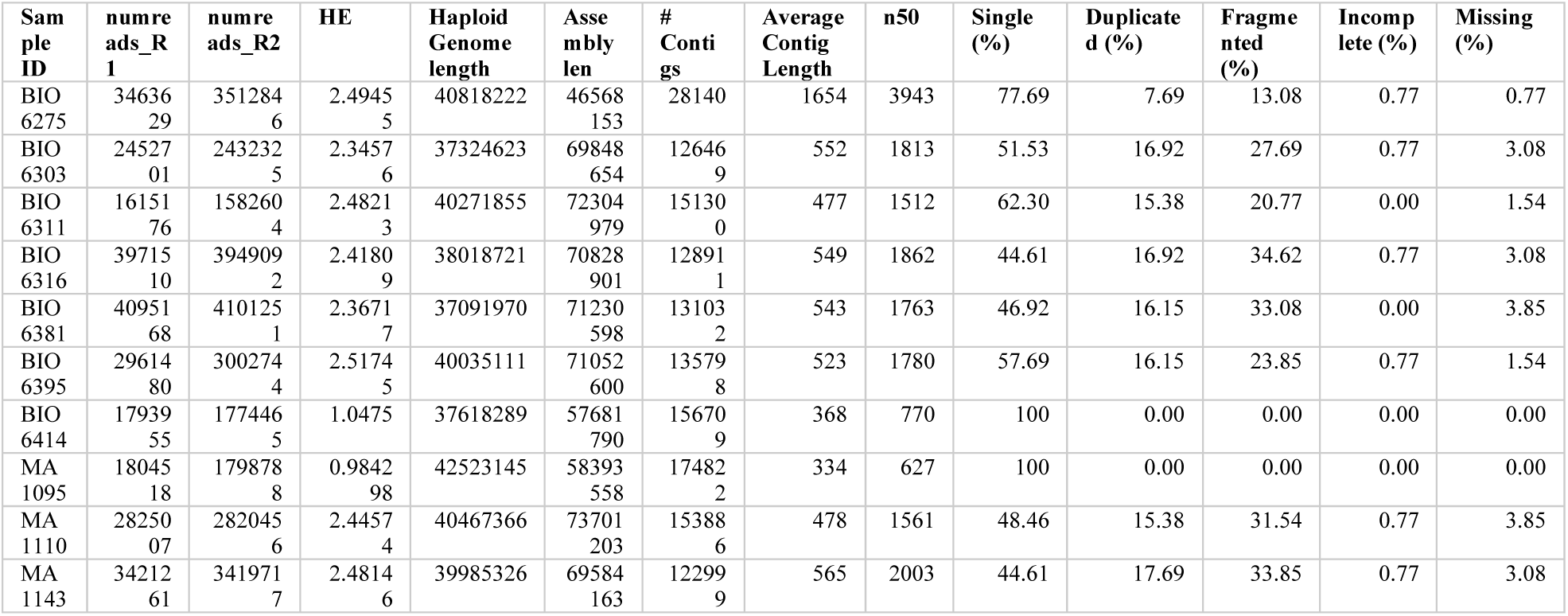
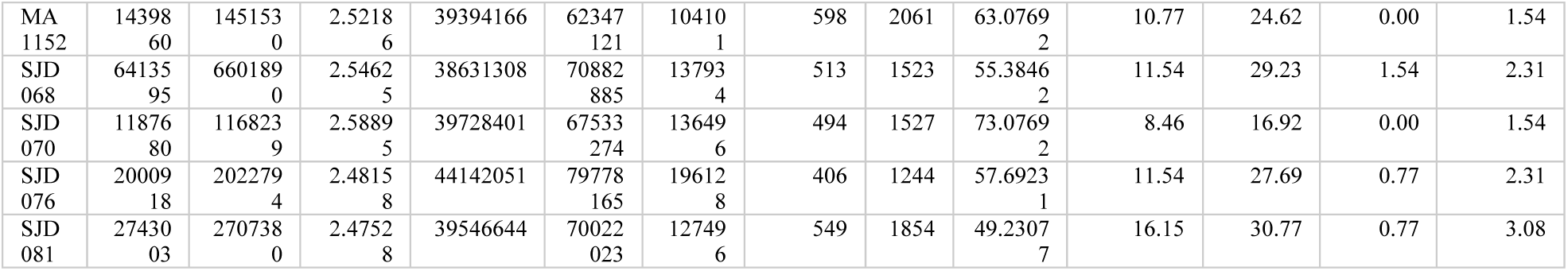
Genome assembly statistics.

| Sam<br>ple<br>ID | numre<br>ads_R<br>1 | numre<br>ads_R2 | HE | Haploid<br>Genome<br>length | Asse<br>mbly<br>len | #<br>Conti<br>gs | Average<br>Contig<br>Length | n50 | Single<br>(%) | Duplicate<br>d (%) | Frage<br>mented<br>(%) | Incomp<br>lete (%) | Missing<br>(%) |
| --- | --- | --- | --- | --- | --- | --- | --- | --- | --- | --- | --- | --- | --- |
| BIO<br>6275 | 34636<br>29 | 351284<br>6 | 2.4945<br>5 | 40818222 | 46568<br>153 | 28140 | 1654 | 3943 | 77.69 | 7.69 | 13.08 | 0.77 | 0.77 |
| BIO<br>6303 | 24527<br>01 | 243232<br>5 | 2.3457<br>6 | 37324623 | 69848<br>654 | 12646<br>9 | 552 | 1813 | 51.53 | 16.92 | 27.69 | 0.77 | 3.08 |
| BIO<br>6311 | 16151<br>76 | 158260<br>4 | 2.4821<br>3 | 40271855 | 72304<br>979 | 15130<br>0 | 477 | 1512 | 62.30 | 15.38 | 20.77 | 0.00 | 1.54 |
| BIO<br>6316 | 39715<br>10 | 394909<br>2 | 2.4180<br>9 | 38018721 | 70828<br>901 | 12891<br>1 | 549 | 1862 | 44.61 | 16.92 | 34.62 | 0.77 | 3.08 |
| BIO<br>6381 | 40951<br>68 | 410125<br>1 | 2.3671<br>7 | 37091970 | 71230<br>598 | 13103<br>2 | 543 | 1763 | 46.92 | 16.15 | 33.08 | 0.00 | 3.85 |
| BIO<br>6395 | 29614<br>80 | 300274<br>4 | 2.5174<br>5 | 40035111 | 71052<br>600 | 13579<br>8 | 523 | 1780 | 57.69 | 16.15 | 23.85 | 0.77 | 1.54 |
| BIO<br>6414 | 17939<br>55 | 177446<br>5 | 1.0475 | 37618289 | 57681<br>790 | 15670<br>9 | 368 | 770 | 100 | 0.00 | 0.00 | 0.00 | 0.00 |
| MA<br>1095 | 18045<br>18 | 179878<br>8 | 0.9842<br>98 | 42523145 | 58393<br>558 | 17482<br>2 | 334 | 627 | 100 | 0.00 | 0.00 | 0.00 | 0.00 |
| MA<br>1110 | 28250<br>07 | 282045<br>6 | 2.4457<br>4 | 40467366 | 73701<br>203 | 15388<br>6 | 478 | 1561 | 48.46 | 15.38 | 31.54 | 0.77 | 3.85 |
| MA<br>1143 | 34212<br>61 | 341971<br>7 | 2.4814<br>6 | 39985326 | 69584<br>163 | 12299<br>9 | 565 | 2003 | 44.61 | 17.69 | 33.85 | 0.77 | 3.08 |
| MA<br>1152 | 14398<br>60 | 145153<br>0 | 2.5218<br>6 | 39394166 | 62347<br>121 | 10410<br>1 | 598 | 2061 | 63.0769<br>2 | 10.77 | 24.62 | 0.00 | 1.54 |
| SJD<br>068 | 64135<br>95 | 660189<br>0 | 2.5462<br>5 | 38631308 | 70882<br>885 | 13793<br>4 | 513 | 1523 | 55.3846<br>2 | 11.54 | 29.23 | 1.54 | 2.31 |
| SJD<br>070 | 11876<br>80 | 116823<br>9 | 2.5889<br>5 | 39728401 | 67533<br>274 | 13649<br>6 | 494 | 1527 | 73.0769<br>2 | 8.46 | 16.92 | 0.00 | 1.54 |
| SJD<br>076 | 20009<br>18 | 202279<br>4 | 2.4815<br>8 | 44142051 | 79778<br>165 | 19612<br>8 | 406 | 1244 | 57.6923<br>1 | 11.54 | 27.69 | 0.77 | 2.31 |
| SJD<br>081 | 27430<br>03 | 270738<br>0 | 2.4752<br>8 | 39546644 | 70022<br>023 | 12749<br>6 | 549 | 1854 | 49.2307<br>7 | 16.15 | 30.77 | 0.77 | 3.08 |

**Supplemental Table 2:**
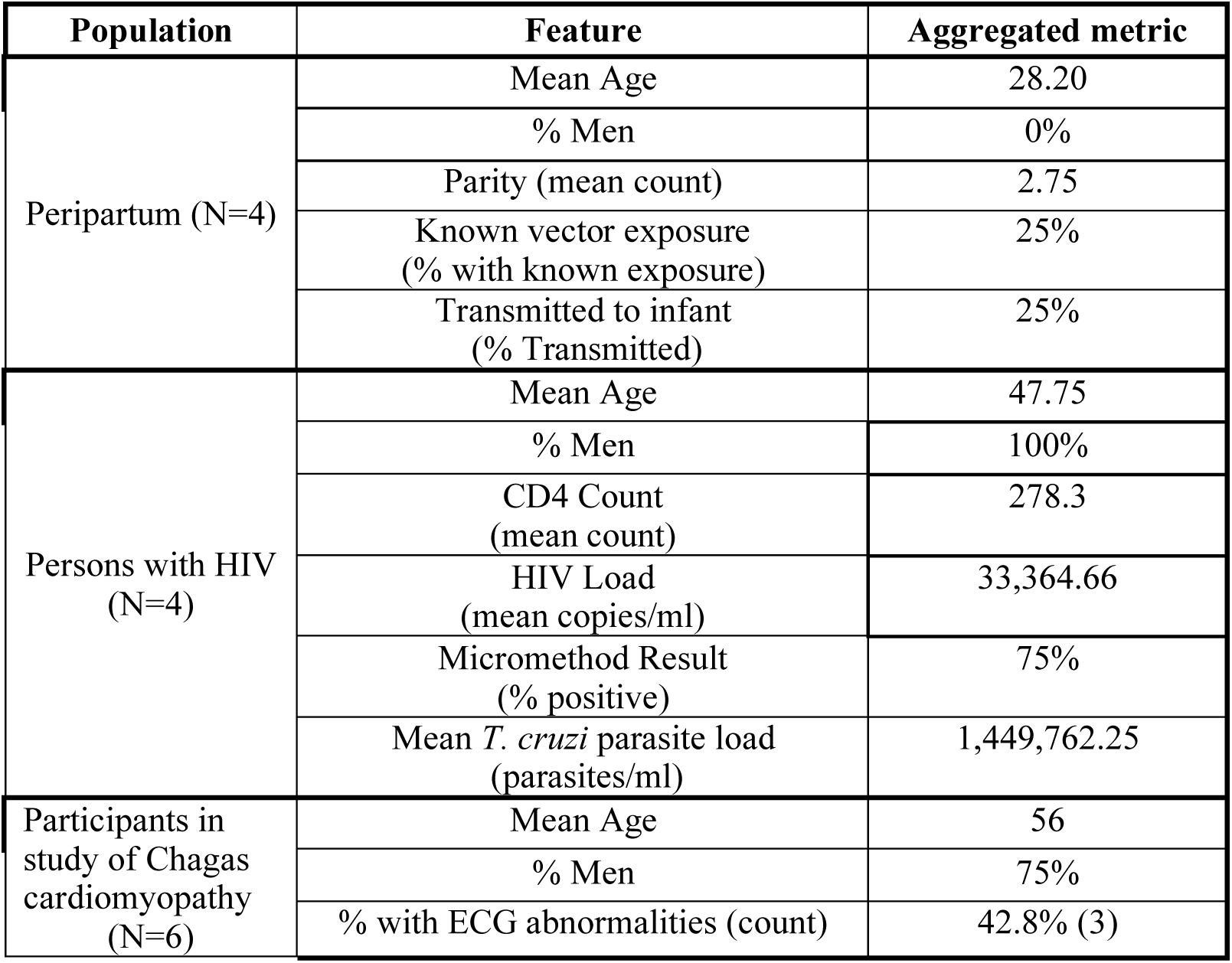
aggregated characteristics of study population

## Notes

### Competing Interest Statement

The authors have declared no competing interest.

