## Supplemental Figures for "Clinical *Trypanosoma cruzi* isolates share a common antigen repertoire that is absent from culture adapted strains"

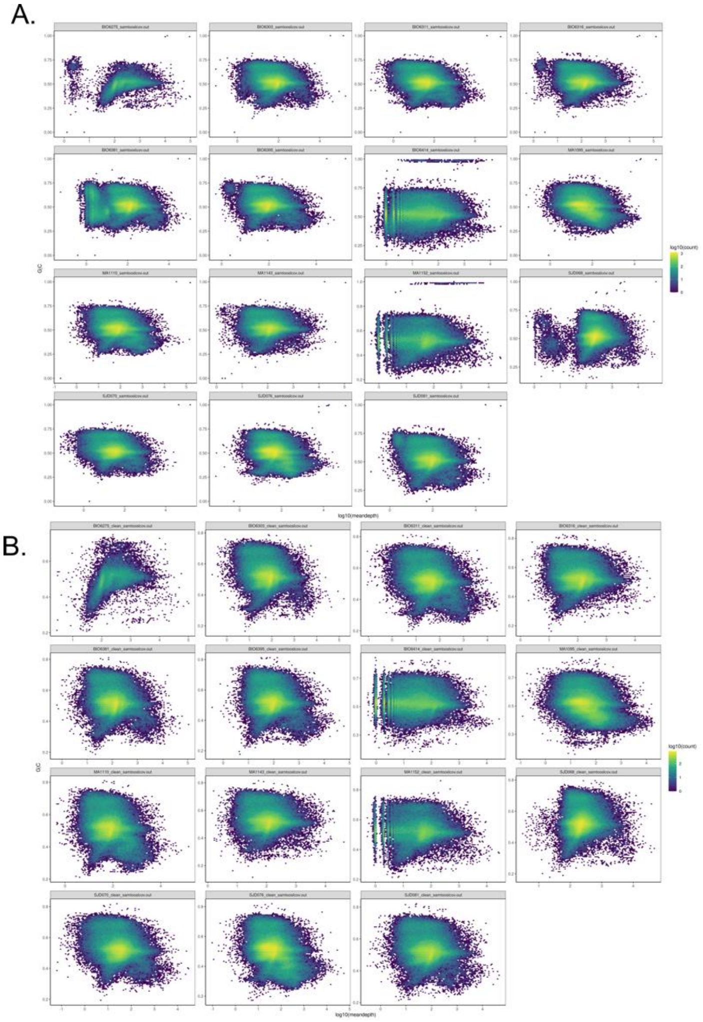


**Supplemental Figure 1**: 2D histograms of GC percent and contig coverage for clinical isolate assemblies before (A) and after (B) cleaning contaminating DNA and kDNA

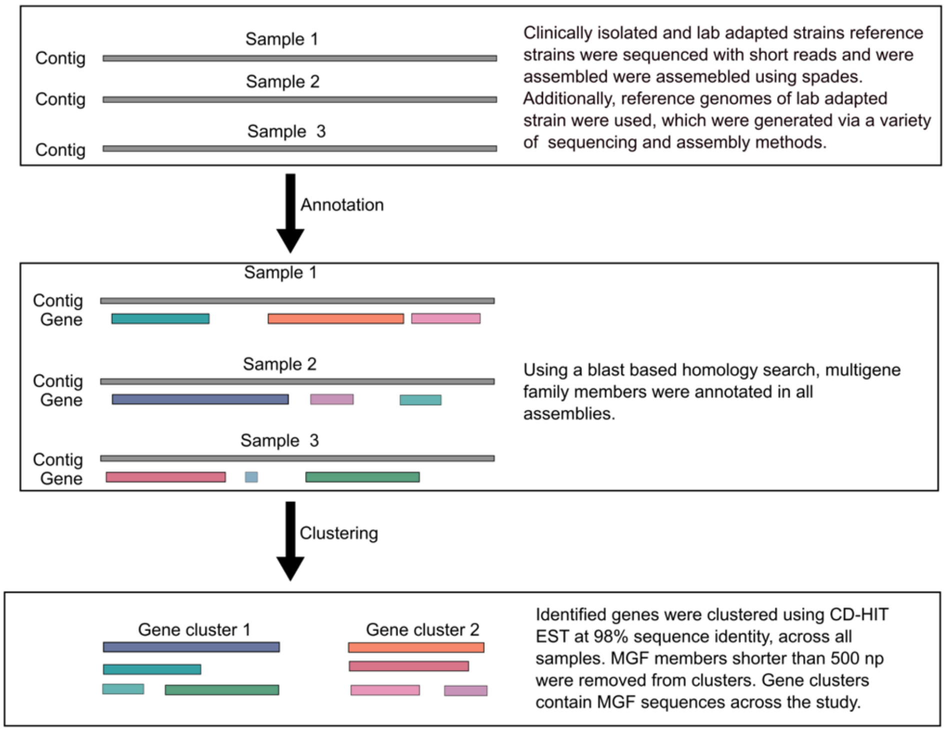


**Supplemental Figure 2**: Schematic representation of gene clustering strategy for multigene family members

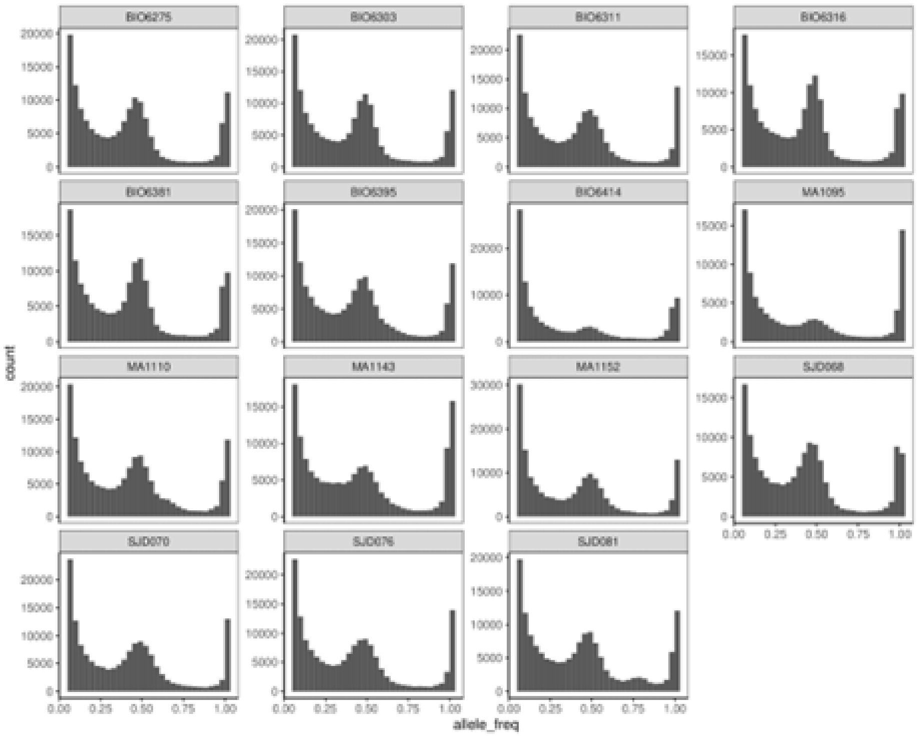


**Supplemental Figure 3**: Distribution of allele frequency for clinically isolated strains against the CL Brener Esmeraldo-like reference genome. Isolates BIO6414 and MA1095 have smaller peaks at 0.5, due to their non-hybrid nature, and lower prevalence of heterozygous alleles.

| Sample ID | numreads_R1 | numreads_R2 | HE | Haploid Genome length | Assembly len | # Contigs | Average Contig Length | n50 | Single (%) | Duplicated (%) | Fragmented (%) | Incomplete (%) | Missing (%) |
|---|---|---|---|---|---|---|---|---|---|---|---|---|---|
| BIO6275 | 3463629 | 3512846 | 2.49455 | 40818222 | 46568153 | 28140 | 1654 | 3943 | 77.69 | 7.69 | 13.08 | 0.77 | 0.77 |
| BIO6303 | 2452701 | 2432325 | 2.34576 | 37324623 | 69848654 | 126469 | 552 | 1813 | 51.53 | 16.92 | 27.69 | 0.77 | 3.08 |
| BIO6311 | 1615176 | 1582604 | 2.48213 | 40271855 | 72304979 | 151300 | 477 | 1512 | 62.30 | 15.38 | 20.77 | 0.00 | 1.54 |
| BIO6316 | 3971510 | 3949092 | 2.41809 | 38018721 | 70828901 | 128911 | 549 | 1862 | 44.61 | 16.92 | 34.62 | 0.77 | 3.08 |
| BIO6381 | 4095168 | 4101251 | 2.36717 | 37091970 | 71230598 | 131032 | 543 | 1763 | 46.92 | 16.15 | 33.08 | 0.00 | 3.85 |
| BIO6395 | 2961480 | 3002744 | 2.51745 | 40035111 | 71052600 | 135798 | 523 | 1780 | 57.69 | 16.15 | 23.85 | 0.77 | 1.54 |
| BIO6414 | 1793955 | 1774465 | 1.0475 | 37618289 | 57681790 | 156709 | 368 | 770 | 100 | 0.00 | 0.00 | 0.00 | 0.00 |
| MA1095 | 1804518 | 1798788 | 0.984298 | 42523145 | 58393558 | 174822 | 334 | 627 | 100 | 0.00 | 0.00 | 0.00 | 0.00 |
| MA1110 | 2825007 | 2820456 | 2.44574 | 40467366 | 73701203 | 153886 | 478 | 1561 | 48.46 | 15.38 | 31.54 | 0.77 | 3.85 |
| MA1143 | 3421261 | 3419717 | 2.48146 | 39985326 | 69584163 | 122999 | 565 | 2003 | 44.61 | 17.69 | 33.85 | 0.77 | 3.08 |
| MA1152 | 1439860 | 1451530 | 2.52186 | 39394166 | 62347121 | 104101 | 598 | 2061 | 63.07692 | 10.77 | 24.62 | 0.00 | 1.54 |
| SJD068 | 6413595 | 6601890 | 2.54625 | 38631308 | 70882885 | 137934 | 513 | 1523 | 55.38462 | 11.54 | 29.23 | 1.54 | 2.31 |
| SJD070 | 1187680 | 1168239 | 2.58895 | 39728401 | 67533274 | 136496 | 494 | 1527 | 73.07692 | 8.46 | 16.92 | 0.00 | 1.54 |
| SJD076 | 2000918 | 2022794 | 2.48158 | 44142051 | 79778165 | 196128 | 406 | 1244 | 57.69231 | 11.54 | 27.69 | 0.77 | 2.31 |
| SJD081 | 2743003 | 2707380 | 2.47528 | 39546644 | 70022023 | 127496 | 549 | 1854 | 49.23077 | 16.15 | 30.77 | 0.77 | 3.08 |

**Supplemental Table 1**: Genome assembly statistics.

| Population | Feature | Aggregated metric |
|---|---|---|
| Peripartum (N=4) | Mean Age | 28.20 |
| | % Men | 0% |
| | Parity (mean count) | 2.75 |
| | Known vector exposure<br>(% with known exposure) | 25% |
| | Transmitted to infant | 25% |

| | | |
|---|---|---|
| | (% Transmitted) | |
| Persons with HIV (N=4) | Mean Age | 47.75 |
| | % Men | 100% |
| | CD4 Count (mean count) | 278.3 |
| | HIV Load (mean copies/ml) | 33,364.66 |
| | Micromethod Result (% positive) | 75% |
| | Mean *T. cruzi* parasite load (parasites/ml) | 1,449,762.25 |
| Participants in study of Chagas cardiomyopathy (N=6) | Mean Age | 56 |
| | % Men | 75% |
| | % with ECG abnormalities (count) | 42.8% (3) |

**Supplemental Table 2**: aggregated characteristics of study population

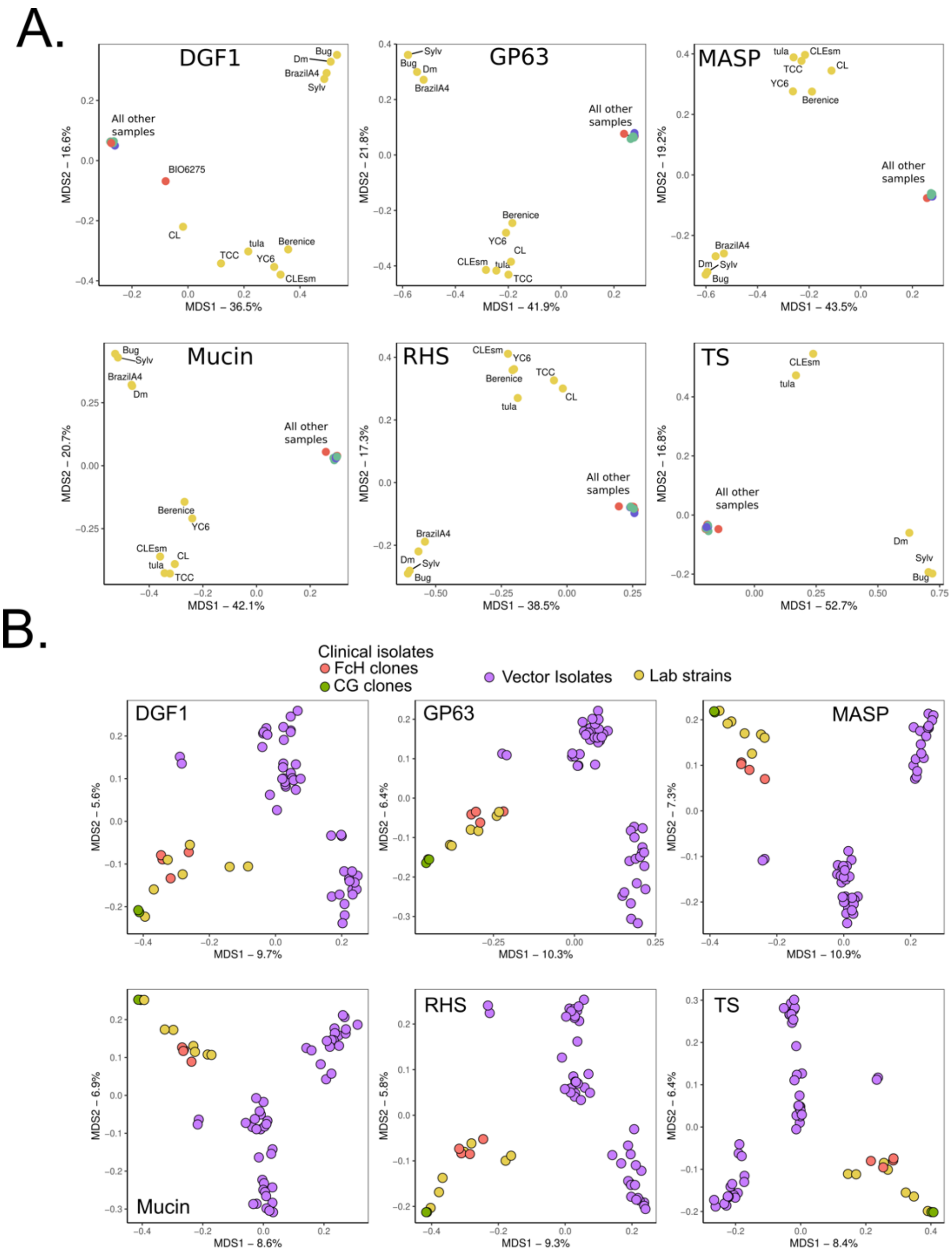


**Supplemental Figure 4**: A) MGF repertoire separation including Tc1 lab reference strains. DTU separation is more pronounced but note that due to the lack of clinically isolated specimens in the current study some separation may be largely DTU driven. B) MGF repertoire separation of TcI parasites alone, including Colombian clinically isolated parasites from Talavera-Lopez et al (FcH and CG, red and green respectively) and vector isolates from Schwabl et al. (purple). FcH and CG are two individuals for which there were four and two parasite genomes cloned from the patient samples

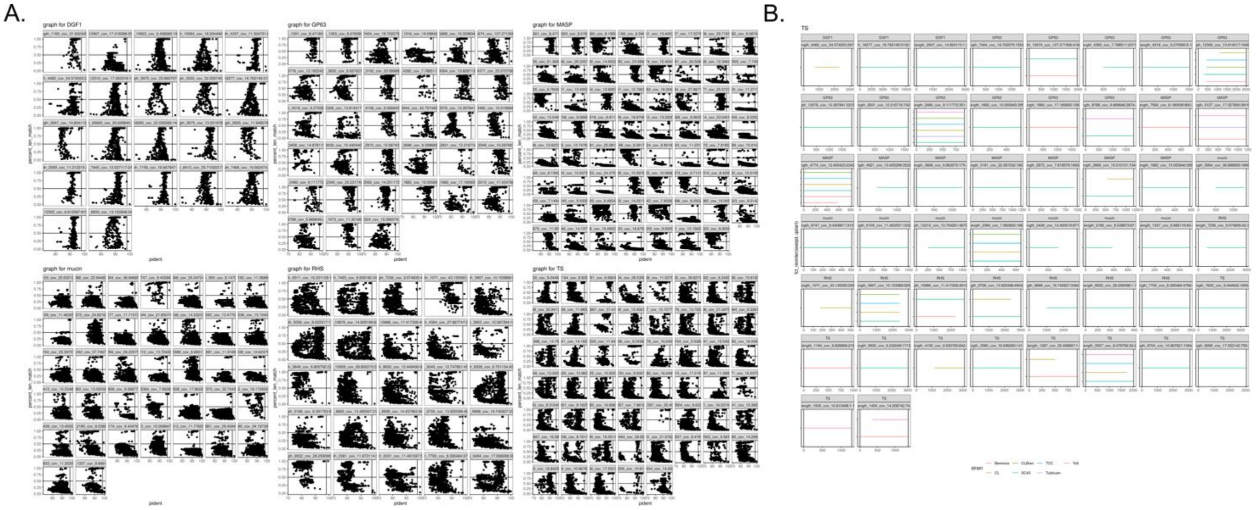


**Supplemental Figure 3:** Confirming the loss of shared MGFs among lab adapted strains by BLASTN search of lost representative sequences within whole genomes of TcII, TcV and TcVI strains. **A)** Plot of all BLASTN hits in reference genomes for each lost MGF encoding gene, across all gene families. X axis represents the percent identity of the BLASTN hit, and Y axis represents the percent of the total MGF that is covered by the BLASTN homology found in the reference genome. The horizontal line is at 50% query coverage, and the vertical line is at 98% BLASTN sequence identity. **B)** Each MGF member that passes the above thresholds. Each gene is a facet, colored by the reference genome the gene was found in. X axis is the length of the gene, with black bars demarcating that start and end, and line segments of each reference genome representing the position where the hit occurred. These are the genes that were removed from the set of shared clinical MGF members that were lost in the lab adapted strains.
